# Design of Vaccine Trials during Outbreaks with and without a Delayed Vaccination Comparator

**DOI:** 10.1101/074088

**Authors:** Natalie E. Dean, M. Elizabeth Halloran, Ira M. Longini

## Abstract

Conducting vaccine efficacy trials during outbreaks of emerging pathogens poses particular challenges. The ‘Ebola ça suffit’ trial in Guinea used a novel ring vaccination cluster randomized design to target populations at highest risk of infection. Another key feature of the trial was the use of a delayed vaccination arm as a comparator, in which clusters were randomized to immediate vaccination or vaccination 21 days later. This approach, chosen to improve ethical acceptability of the trial, complicates the statistical analysis as participants in the comparison arm are eventually protected by vaccine. Furthermore, for infectious diseases, we observe time of illness onset and not time of infection, and we may not know the time required for the vaccinee to develop a protective immune response. As a result, including events observed shortly after vaccination may bias the per protocol estimate of vaccine efficacy. We provide a framework for approximating the bias and power of any given per protocol analysis period as functions of the background infection hazard rate, disease incubation period, and vaccine immune response. We use this framework to provide recommendations for designing standard vaccine efficacy trials and trials with a delayed vaccination comparator. Briefly, narrower analysis periods within the correct window can minimize or eliminate bias but may suffer from reduced power. Designs should be reasonably robust to misspecification of the incubation period and time to develop a vaccine immune response.

## 1 Background

Evaluating the efficacy of a vaccine candidate during a public health emergency brings unique challenges. Trials in resource-limited settings may face severe logistical constraints; transmission can be highly localized and hard to predict; furthermore, the ethics of a trial in the face of an emergency are complex. Innovative designs can address some of these challenges. During the 2013-2016 West African Ebola epidemic, the ‘Ebola ¸ca suffit’ ring vaccination trial in Guinea randomized clusters of contacts and contacts of contacts of confirmed Ebola virus disease cases to immediate or delayed (after 21 days) vaccination [Ebola c¸a Suffit Ring Vaccination Trial Consortium, 2015]. The trial used the delayed vaccination arm as a comparator to estimate vaccine efficacy (VE).

When analyzing data from a vaccine trial with or without a delayed vaccination arm, infection times are unknown and only illness (or symptom) onset times are observable. The time between infection and illness onset, known as the incubation period, is an unobserved random variable. Furthermore, there is a delay between vaccination and the development of a robust immune response which we refer to as the vaccine ramp-up period. As a result, individuals with illness onsets occurring shortly after vaccination were likely infected prior to vaccination (incubating cases) and/or were infected when the immune response had not fully developed. Including these cases in the per protocol analysis can bias the estimated VE towards the null [Horne et al., 2000]. In response, the analysis period for vaccine trials often starts after some delay greater than the maximum incubation period plus the maximum vaccine ramp-up period. For example, the per protocol analysis of the RTS,S malaria vaccine was restricted to cases occurring 14 or more days after the third dose [RTSS Clinical Trials Partnership, 2015]. Unfortunately, this approach can negatively impact power. In the RV144 HIV vaccine efficacy trial, the per protocol analysis (seronegative at 26 week visit and followed protocol) was statistically non-significant because it excluded about 25% of study participants and 31% of infections, whereas the modified intent to treat analysis (seronegative at baseline visit) was significant at the 0.05 level [Gilbert et al., 2011]. No formal guidance exists on how to identify the optimal analysis period for vaccine trials.

A second key design issue is that the comparator arm may receive delayed vaccination for ethical reasons, as was used in the ‘Ebola ça suffit’ trial [Henao-Restrepo et al., 2015]. This has a few benefits, including that all trial participants receive the vaccine while they are still at risk. Participants may also be more likely to consent if they are offered delayed vaccination as compared to placebo or vaccination for a different disease. Finally, trial partners may be more likely to approve a protocol with this component. The use of delayed vaccination necessarily decreases study power as it restricts the period when the comparator arm is unprotected by vaccine and thus VE is estimable. This reduction in power as compared to a standard parallel design is well-recognized for stepped wedge trials, in which all clusters start in the control arm and then receive the intervention in a randomized order [Hussey and Hughes, 2007, Bellan et al., 2015]. Delayed vaccination is related to this approach, and it can be viewed as a one-way crossover trial. As the use of a delayed vaccination comparator arm outside of stepped wedge trials is novel, we provide guidance on how to select this vaccination delay. Furthermore, we make recommendations on how to conduct the analysis in the presence of a delayed vaccination arm, including when to start including cases in the primary analysis and, importantly, when to stop including cases.

In this paper, we describe a model for the hazard rates of illness onset times in a vaccine trial with and without delayed vaccination. We present closed-form approximations for apparent VE and power to detect a significant vaccine effect for any given analysis period. We describe a framework for selecting the optimal analysis period start in terms of maximizing power and minimizing bias in trials with a placebo or unrelated vaccine control arm. Next, we consider trials using delayed vaccination and provide recommendations on selecting this delay and conducting the analysis. We describe the principles in the context of the Guinea Ebola vaccine trial.

## 2 Description of the model

We describe a model for the observable illness onset hazard rate as a function of the unobservable infection hazard rate, time-dependent vaccine protection, and disease incubation period. Figure 1 describes the key variables.

**Figure 1:**
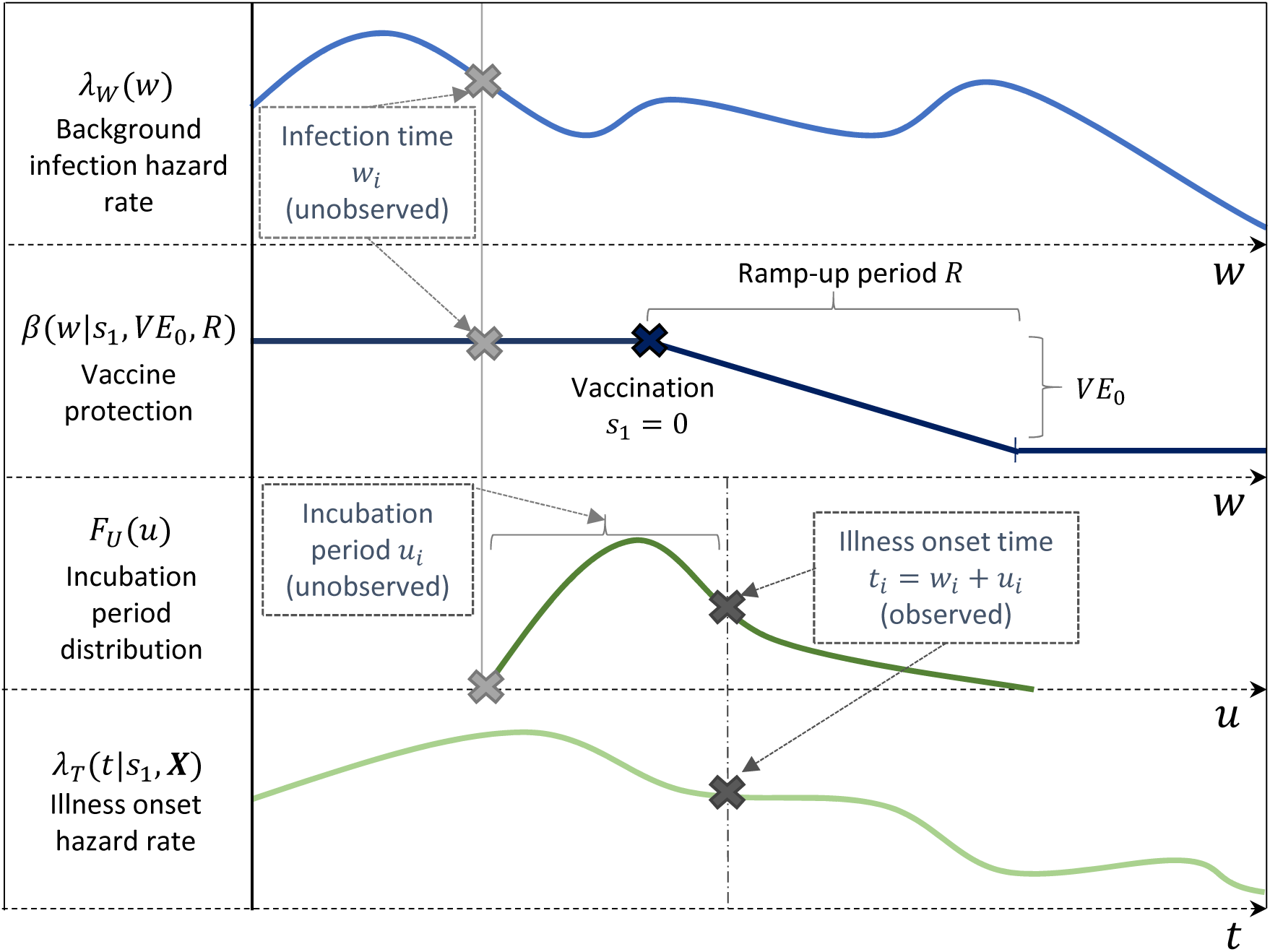
Hypothetical infected individual *i* in Arm 1 (intervention arm). [*First row*] The unobserved infection time *w*_*i*_ occurs with background infection hazard rate *λ*_*W*_ (*w*_*i*_). [*Second row*] At infection time *w*_*i*_, individual *i* has vaccine protection *β* (*w_i_|s*_1_, *VE*_0_, *R*) which has no impact before the individual is vaccinated and reduces the hazard by *VE*_0_ once the vaccine is fully protective. [*Third row*] The observed illness onset time *t*_*i*_ is the sum of the unobserved infection time *w*_*i*_ and incubation period *u*_*i*_, where *u*_*i*_ is drawn from the incubation period distribution *F*_*U*_ (*u*_*i*_). [*Last row*] Only illness onsets and the illness onset hazard rate *λ*_*T*_ (*t|s*_1_, **X**) are observable and estimable from the data, letting **X** be the set of parameters comprised of *VE*_0_, *R, F*_*U*_ (*•*) and *λ*_*W*_ (*•*).

The background infection hazard rate, defined as the rate at which a fully susceptible unvaccinated person who is exposed becomes infected, is a time-dependent function *λ*_*W*_ (*w*) of infection times *W* = *w*. In the absence of vaccination, the actual infection hazard rate is equal to the background infection hazard rate. As the immune response develops, the background infection hazard rate is multiplicatively reduced by the vaccine effect (partial or leaky vaccine protection) [Halloran et al., 1991]. The “vaccine ramp-up” period is the time from vaccination to maximum efficacy *VE*_0_ and has duration *R*. During the ramp-up period, vaccine protection is assumed to be linearly increasing, though this could be readily modified. *β* (*w|s_j_, V E*_0_, *R*) below describes the protective vaccine effect at infection time *w* in Arm *j* vaccinated on day *s*_*j*_.

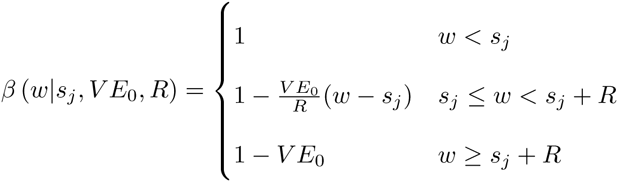

In a standard vaccine trial, the intervention arm (Arm 1) is vaccinated on day *s*_1_ = 0, and the comparator arm (Arm 0) is never vaccinated, *s*_0_ = *∞*. In a trial with a delayed vaccination arm, the immediate arm (Arm 1) is vaccinated on day *s*_1_ = 0, and the delayed arm (Arm 0) is vaccinated on day *s*_0_ = *b* where *b* is the vaccination delay. Where trial participants are vaccinated on different days (e.g. stepped rollout as in ring vaccination), the time scale might be standardized so that day 0 is the date of randomization. In a ring vaccination trial, for example, the background infection hazard rate can be interpreted as the the infection hazard rate in a population with a recent case that triggered the definition of a ring, assuming that the process of contact tracing and enrollment into the trial will have a similar impact on the infection hazard rate across rings.

Given infection at time *W* = *w*, the incubation period for an infected individual is a non-negative random variable *U* = *u* that follows some continuous distribution with probability distribution function *f*_*U*_(⋅), cumulative distribution function *F*_*U*_(⋅), median *u*_0.50_, 90th percentile *u*_0.90_, and 99.9th percentile *u*_0.999_. Examples of distributions are *U ∼ Unif* (0, 10) or *U ∼ Gamma*(*shape* = 6, *scale* = 1). Illness onset time is thus a random variable *T* = *t*, where individual *i* develops illness at time *t*_*i*_ = *w*_*i*_ + *u*_*i*_. We assume that *t*_*i*_ is observable for any individual, though *u*_*i*_ and *w*_*i*_ are not. There could be exceptions, such as a known point exposure, but we do not consider these here.

We let **X** be the set of parameters comprised of *VE*_0_, *R, F*_*U*_ (⋅) and *λ*_*W*_ (⋅). Integrating over all possible combinations of infection times and incubation periods, the illness onset hazard rate 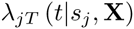 at a particular time *t* in Arm *j* ∈ {0, 1} has the following form:

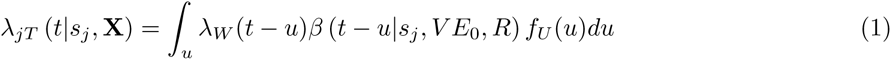

The illness onset hazard rate simplifies nicely in a few settings. When the background infection hazard rate is constant (*λ*_*W*_ (*w*) = *λ_W_ ∀w*) and there is no ramp-up period (*R* = 0), the illness onset hazard rate at time *t* in Arm 1 is as follows:

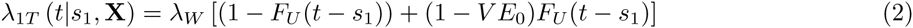

Equation (2) suggests that, when the background infection hazard rate is constant, for a given time *t*, we consider all combinations of incubation periods *u* and infection times *w* = *t* − *u*. Infections after vaccination are downweighted by *VE*_0_, making infections prior to vaccination relatively more likely to occur.

Returning to the general setting, the apparent VE at time *t*, denoted *VE*_*T*_ (*t*), is equal to one minus the apparent illness onset hazard ratio, written below:

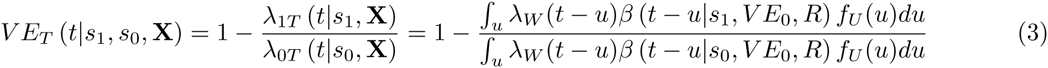

For notational simplicity, we omit the conditioning arguments in the following sections.

## 3 Illness onset hazard rates and apparent VE

We use the equations in Section 2 to model hypothetical vaccine trials. To support meaningful comparisons, we primarily use a single “standard” setting in our examples modeled after Ebola virus disease where the disease incubation period is gamma distributed with a mean of 6 days and scale parameter of 1. The vaccine is fast-acting with a four day vaccine ramp-up period (*R* = 4) before maximal VE of *VE*_0_ = 90% is achieved. The biggest expected challenge is a low event rate and potentially decreasing background infection hazard rate as disease containment procedures are implemented.

Figure 2 depicts the illness onset hazard rates in each arm, *λ*_1*T*_(*t*) and *λ*_0*T*_(*t*), and apparent VE, *VE*_*T*_ (*t*), as determined by Equations (1) and (3), respectively. In this standard setting example, Arm 0 is a control arm (no vaccination), and the background infection hazard rate is constant and low (*λ*_*W*_ = 0.001, resulting in an Arm 0 attack rate *≈*3% over 30 days). (An example with decreasing background infection hazard rate is provided in Figure S8 in the Supplementary Materials.) We see that the illness onset hazard rate in Arm 1 drops sharply, though not immediately, after vaccination on day *s*_1_ = 0. Apparent VE increases from 0 to *VE*_0_ = 90% and then stabilizes. The duration of time until stabilization depends on the incubation period and ramp-up distributions; here, stabilization begins shortly before *R* + *u*_0.999_.

**Figure 2:**
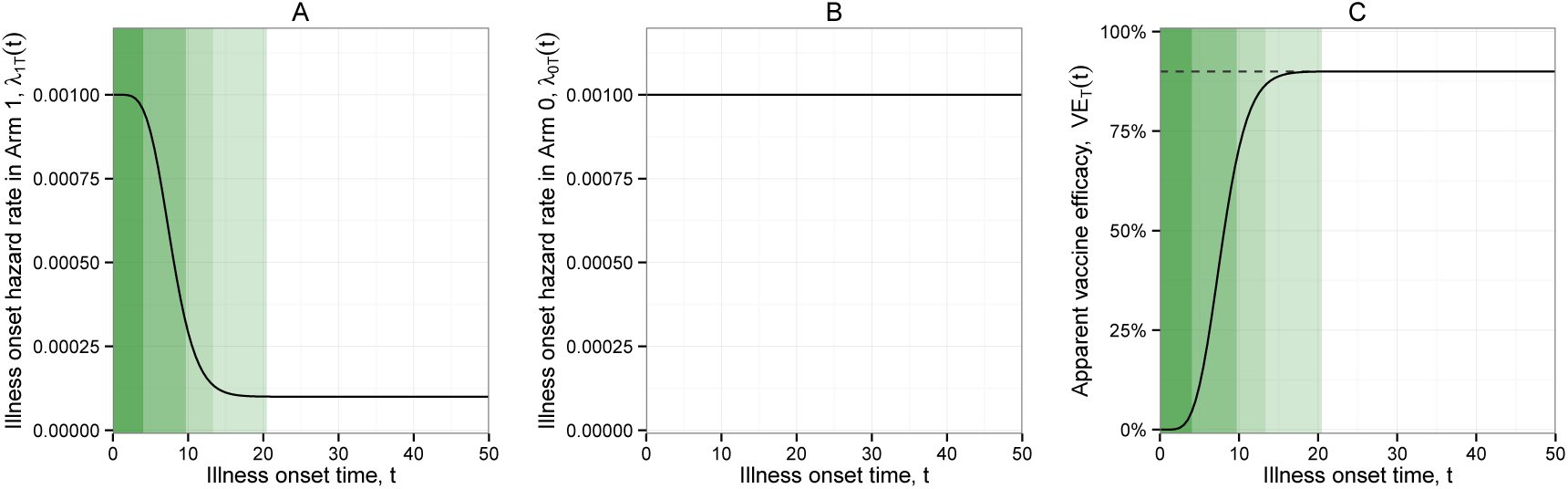
Arm 1 (intervention arm) vaccinated on day *s*_1_ = 0; Arm 0 (comparator arm) never vaccinated (*b* = ∞). Constant background infection hazard rate *λ*_*W*_ = 0.001. Standard setting. For this and all following figures, green shading regions identify key time points in Arm 1, from darkest to lightest, (i) *s*_1_ to *s*_1_ + *R*, (ii) to *s*_1_ + *R* + *u*_0.50_, (iii) to *s*_1_ + *R* + *u*_0.90_, and (iv) to *s*_1_ + *R* + *u*_0.999_. (A) Illness onset hazard rate in Arm 1 as a function of time. (B) Illness onset hazard rate in Arm 0 as a function of time. (C) Apparent VE (1 - illness onset hazard ratio) as a function of time, where the dotted line indicates *VE*_0_.

Figure 3 depicts the illness onset hazard rates in each arm and apparent VE as a function of time when a 21 day vaccination delay (*b* = 21) is used. A constant background infection hazard rate is assumed. (An example with decreasing background infection hazard rate is provided in Figure S9 in the Supplementary Materials.) The illness onset hazard rate in Arm 0 resembles that of Arm 1 shifted by *b* days. Apparent VE, *VE*_*T*_ (*t*), briefly maximizes at *VE*_0_ = 90% but returns to 0 shortly after delayed vaccination. Panel C suggests that there is a limited time window in which *VE*_0_ is estimable; at illness onset times before and after this window, apparent *VE*_*T*_ (*t*) can be highly attenuated.

**Figure 3:**
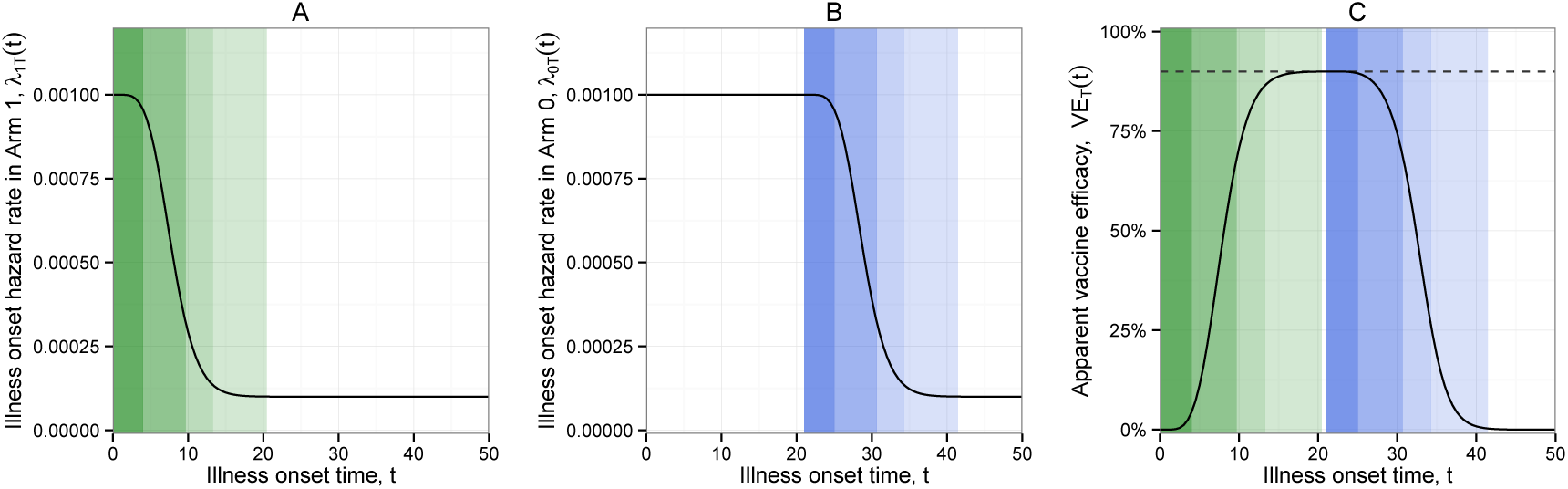
Arm 1 (intervention arm) vaccinated on day *s*_1_ = 0; Arm 0 (comparator arm) vaccinated on day *s*_1_ = 21 (*b* = 21). Constant background infection hazard rate *λ*_*W*_ = 0.001. Standard setting. For this and all following figures, blue shading regions identify key time points in Arm 0, from darkest to lightest, (i) *s*_0_ to *s*_0_ + *R*, (ii) to *s*_0_ + *R* + *u*_0.50_, (iii) to *s*_0_ + *R* + *u*_0.90_, and (iv) to *s*_0_ + *R* + *u*_0.999_. (A) Illness onset hazard rate in Arm 1 as a function of time. (B) Illness onset hazard rate in Arm 0 as a function of time. (C) Apparent VE as a function of time, where the dotted line indicates *VE*_0_.

## 4 Trial analysis framework

As demonstrated in Section 3, apparent VE can be very biased at particular illness onset times, especially when delayed vaccination is used. A key design choice is then which illness onset times should contribute to the per protocol analysis. A simple and commonly used approach is to apply hard cutpoints to create an analysis period (see Figure 4). We use *d* to denote the analysis period start, which is the earliest illness onset day, relative to the date of randomization, included in the per protocol analysis, and we use *c* to denote the width of the analysis period in days. Events occurring before *d* are excluded, and observations are right-censored at time *d* + *c*. The analysis period is identical in both trial arms to maintain comparability; different treatment of the two arms could induce bias [Camacho et al., 2015].

**Figure 4:**
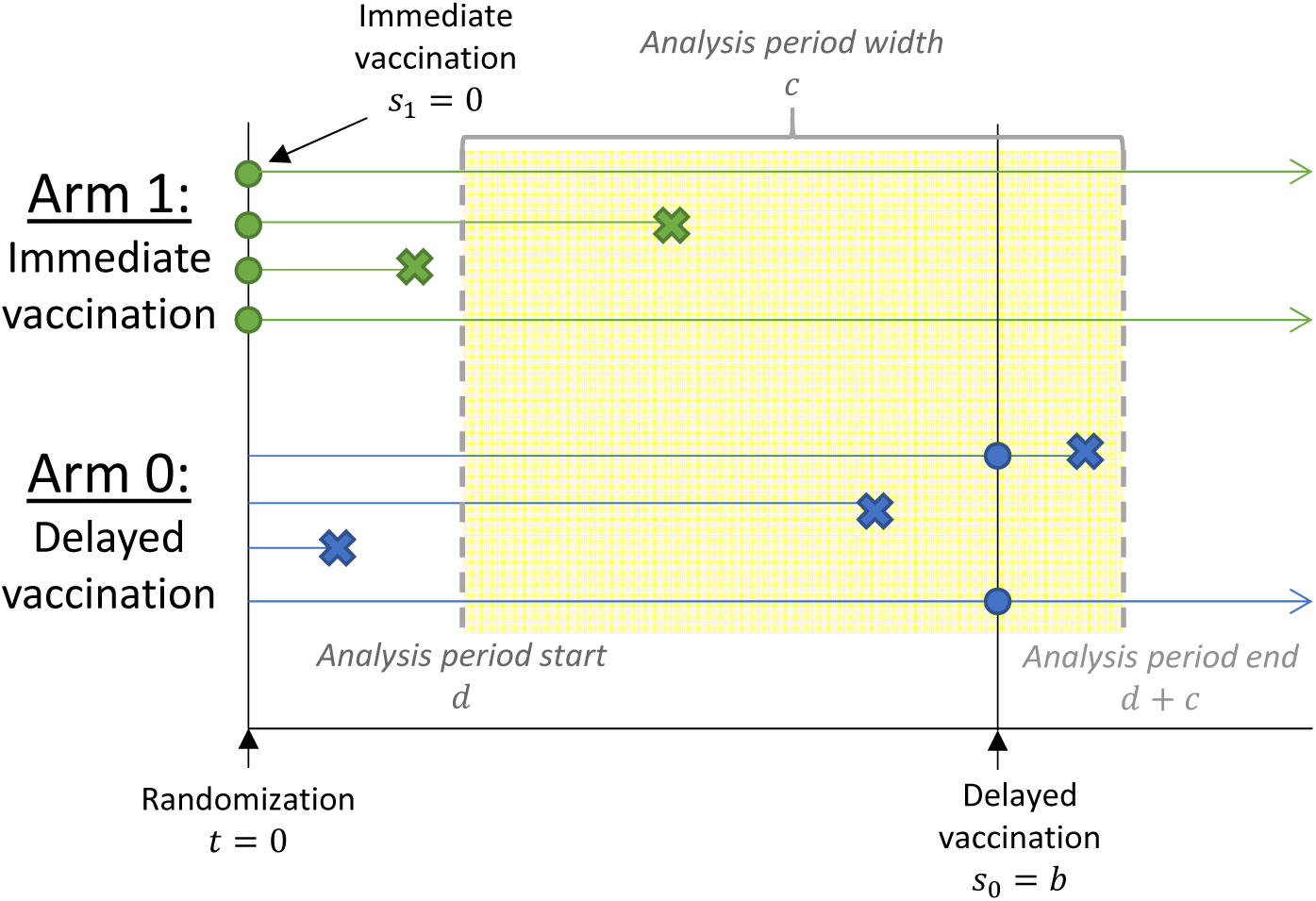
Hypothetical vaccine trial. Green dots indicate immediate vaccination (*s*_1_ = 0) in Arm 1, and blue dots indicate delayed vaccination (*s*_0_ = *b <*) in Arm 0. Xs indicate illness onset dates in the respective trial arms. The yellow shaded area marks the analysis period [*d, d* + *c*) such that only events in this period are included. Events occurring before *d* and after *d* + *c* are excluded.

Cox proportional hazards or a piecewise exponential model fit with a log-linear approach can be used to estimate vaccine efficacy if the data are expressed in time-to-event format with times shifted to subtract the analysis period start *d* to prevent immortal time bias before *d*. These models allow for flexibility in the background infection hazard rate and can accommodate additional covariates (e.g. risk factors for infection). Clustering, if a cluster randomized trial design is used, can be modeled with a shared frailty term.

### 4.1 Bias approximation

In the Cox and piecewise exponential models, illness onset hazard rates *λ*_*T*1_(*t*) and *λ*_*T*0_(*t*) can be time-dependent as long as the hazards are proportional (constant hazard ratio). From Section 3, we see that apparent VE, defined as one minus the hazard ratio, is actually time-varying (“time varying effect”). When we assume proportional hazards but proportional hazards is violated, we estimate an average regression effect [Xu and O’Quigley, 2000]. As no closed-form expression is available for the Cox model, we approximate the expected apparent VE for a given analysis period, 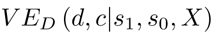, from the ratio of the average illness onset hazard rates observed in [*d*, *d* + *c*) (see Equation (4)).

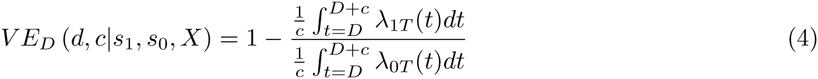

We evaluate the performance of this approximation using a simulation approach in R [R Core Team, 2015] using the survival package [Therneau, 2015] described in the Supplementary Materials. Briefly, approximated VE generally agrees with the simulated average VE within a few percent. This approximation performs best when the background infection hazard rate *λ*_*W*_ (⋅) is highest and/or the sample size per arm *n* is largest. We do not consider clustered data here as clustering does not impact the expected bias.

### 4.2 Power approximation

The power for comparing survival curves under Cox proportional hazards can be approximated by the power of the log rank test statistic to test *H*_0_: *VE*_0_ ≠ 0% [Rosner, 2010] (Chapter 14, Page 784). To estimate power for a given analysis period, 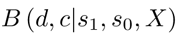, we adapt the standard formula, replacing the true VE, *VE*_0_, with the apparent VE, *VE*_*D*_(*d, c*), and calculating the expected number of events (illness onsets) using the illness onset hazard rates.

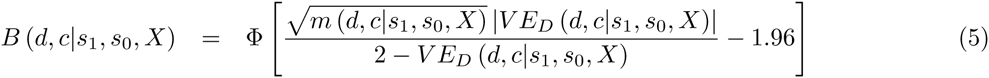

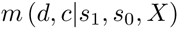 is the expected total number of illness onsets during the analysis period, calculated as *np*_1_ + *np*_0_ where *n* is the sample size per arm and *p*_*j*_ (below) is the probability that a participant does not have an illness onset occurring before time *d* but has an illness onset during the analysis period [*d, d* + *c*) in arm *j* ∈ {0, 1}. Though a time-dependent background infection hazard rate could be used, in practice, one would likely assume a constant rate.

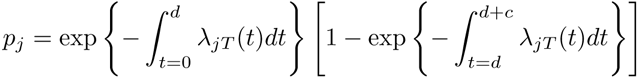

The adequacy of this approximation is evaluated with simulations in the Supplementary materials (see Tables S1, S2, S3, and S4). Generally there is good agreement (*∼*2% absolute difference) between simulated and approximated power, though the approximation tends to slightly overestimate power. The approximation performs best when the sample size is large. The approximation performs poorly when the expected number of events in Arm 1, intervention arm, is small, e.g. less than 5, and can perform poorly when the sample size is small, e.g. *n* = 500.

Equation (5) can be adapted to test *VE*_0_ *>* 30% or some other pre-specified lower bound. Results are presented for individually randomized trials, but this can be modified for cluster designs by solving Equation (5) for the effective sample size *n* and then multiplying the effective sample size by the trial design effect. The design effect is frequently approximated as 1 + (*m −* 1)*ρ* where *ρ* is the intracluster correlation coefficient and *m* is the per-cluster sample size [Ridout et al., 1999].

## 5 Trials with an unvaccinated control arm

After establishing methods for approximating the apparent VE and power for a given trial design (i.e., choice of *d* and *c*), we investigate how to select *d* and *c* in the simple setting of a vaccine trial in which Arm 0 is never vaccinated with the candidate vaccine. In Figure 5, we consider the setting of a fixed analysis period end *d* + *c* and variable analysis period start *d*. This could occur if the budget only supports follow-up for a fixed time from randomization, and there is incentive to maximize the use of this follow-up time. An alternative scenario with fixed analysis period width *c* and variable analysis period start *d* is considered in the Supplementary Materials (see Figures S10 and S11). We see that *d* below *R* + *u*_0.90_ is associated with bias because it includes a period when the vaccine is only partially effective and apparent VE is biased (see Figure 2). Interestingly, the power maximizes at an analysis period start *d* for which there is some bias in the apparent VE. This demonstrates a balance between bias and variance, with an earlier start *d* capturing more events during the period of partial efficacy, thereby increasing power while slightly biasing apparent VE. When the background infection hazard decreases, the power drops off more steeply for larger values of *d* (see Figure S12 in the Supplementary Materials).

**Figure 5:**
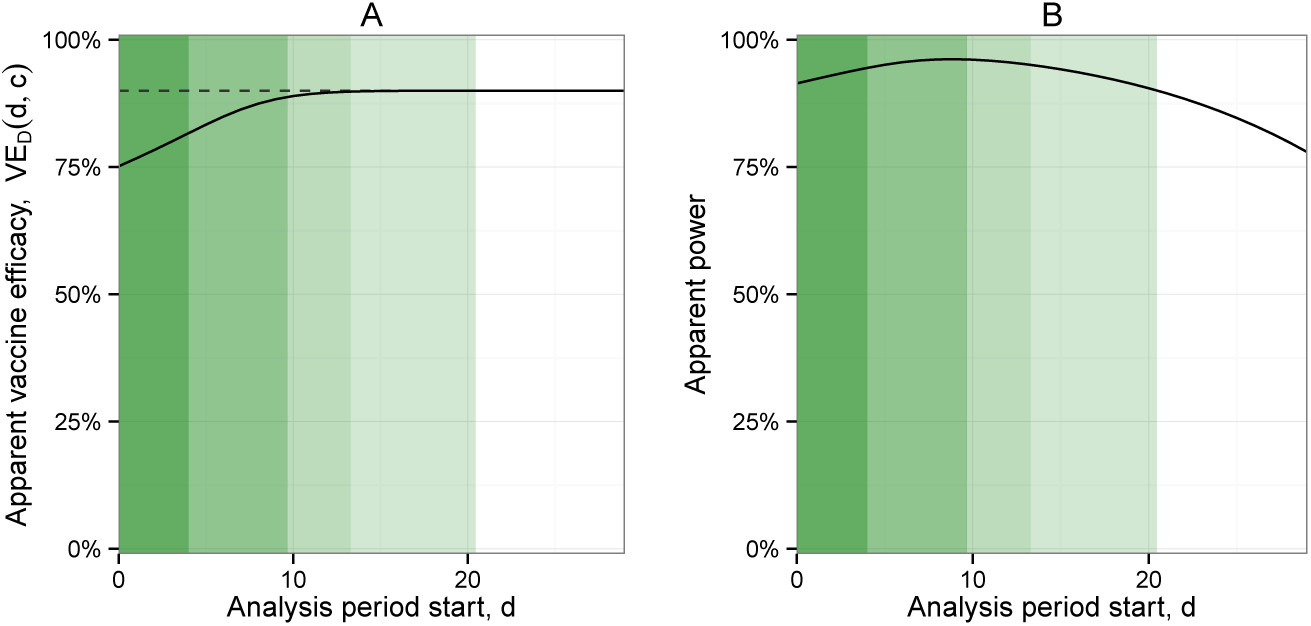
Arm 1 (intervention arm) vaccinated own day *s*_1_ = 0; Arm 0 (comparator arm) never vaccinated (*b* =). Constant background infection hazard rate *λ*_*W*_ = 0.001. Standard setting. Sample size *n* = 500 per arm. (Setting as in Figure 2.) Analysis period [*d,* 50) with width *c* = 50 − *d*; a range of *d* values are considered. (A) Apparent VE to assess bias, where the dotted line indicates *VE*_0_. (B) Apparent power.

We assess the impact of other key factors in the Supplementary Materials, including the incubation period distribution, the ramp-up period, *VE*_0_, and the background infection hazard rate. We summarize the results here.

Longer incubation periods and ramp-up periods shift the optimal *d* but the same general relationships persist. Illness onsets following long incubation periods are most likely to contribute to bias, but the probability of these types of incubation periods is reflected in the distribution’s quantiles, *u*_0.90_ and *u*_0.999_. In this scenario, to minimize bias, an appropriate starting point for the delay could be *R* + *u*_0.90_ or *R* + *u*_0.999_. Examples with different incubation period distributions are provided in Supplementary materials (Figures S13 and S14). For longer ramp-up periods, the power tends to maximize at earlier values of *d* when the ramp-up period is longer because there is a longer period of partial efficacy. If there is little protective effect until the end of the ramp-up period (e.g. ramp-up is not linear), we will need to select a later *d* because an earlier *d* might be prone to more bias (vaccinated group looks similar to unvaccinated group for longer). Conversely, fast-acting vaccines require an earlier *d*. Examples are provided in the Supplementary materials (Figure S18). In an extreme situation, a vaccine with post-exposure prophylactic effects will require the earliest *d* because vaccinees may be protected from disease even if already infected.

The impact of *VE*_0_ is very minimal. The optimal *d* for minimizing bias increases only slightly with increasing *VE*_0_ (see Figure S19 in Supplementary Materials). When *VE*_0_ is highest, the apparent VE experiences the greatest increase from 0; thus, including a period of partial efficacy when the vaccine is highly effective can induce the largest absolute bias. The effect of varying the background infection hazard rate *λ*_*W*_ (*w*) is also minimal. In the constant hazard setting, there is no change in bias, though the mean squared error decreases as the event rate increases. The effect of event rate on power is as expected, with greater power at the higher event rate, but the location of the optimal value of *d* is fairly constant (see Figure S20 in Supplementary Materials).

Given the above results, it is clear that choosing a later analysis period start, *d*, is preferable for minimizing bias between the apparent VE and true *VE*_0_. A safe and commonly used option is to select *d* equal to *R* + *u*_0.999_, the maximum ramp-up period plus the maximum incubation period, though using the 90th or 95th percentile of the incubation period distribution can also give good results. Nonetheless, our results suggest that when maximizing power, there are many settings where we may prefer an earlier value of *d*. Capturing cases during this period of partial vaccine efficacy may provide critical additional events, especially important if the background infection hazard rate is low and/or decreasing. Similarly, an earlier *d* may allow for a wider analysis period *c*, further increasing the event rate.

## 6 Trials with a delayed vaccination arm

Next we consider delayed vaccination of the comparator arm, Arm 0. This design may be adopted to improve ethical acceptability of the trial to partners and participants. Using the framework described above, we investigate how to select an appropriate trial design, defined as the vaccination delay *b*, the analysis start period *d*, and the analysis period width *c*. The choice of *d* follows many of the same principles described in Section 5 and should be based on the expected ramp-up and incubation periods and background infection hazard rate, but the trial results are more sensitive to the choice of *c* when delayed vaccination is used. Issues of power are especially critical because cases must occur during the narrow window before the delayed arm is protected.

A natural starting point is to consider analysis periods that have width equal to the vaccination delay (*c* = *b*). This approach has a nice symmetry because the cutoff (*d*) applied to the immediate arm parallels the cutoff (*b* + *d*) applied to the delayed arm. In practice, if *c* = *b*, there is no value of *d* such that apparent *VE*_*D*_ is unbiased because any analysis period captures either unprotected immediate vaccinees or protected delayed vaccinees. For example, in Figure 3, there is no consecutive 21 day period during which apparent *VE*_*T*_ is unbiased, so an average over this period will also be biased, as shown in Figure 6. Because this approach has a wider analysis period than designs with *c < b*, it typically maximizes power, but power can be highly sensitive to the choice of *d*, with a narrow maximum and steep decline on either end, leaving little tolerance for error in pre-specification. Power can be increased by increasing the sample size (see Figure S22 in Supplementary Materials). These results suggest that, if the primary goal is to reduce bias, we should decouple the vaccination delay *b* and the analysis period width *c*.

**Figure 6:**
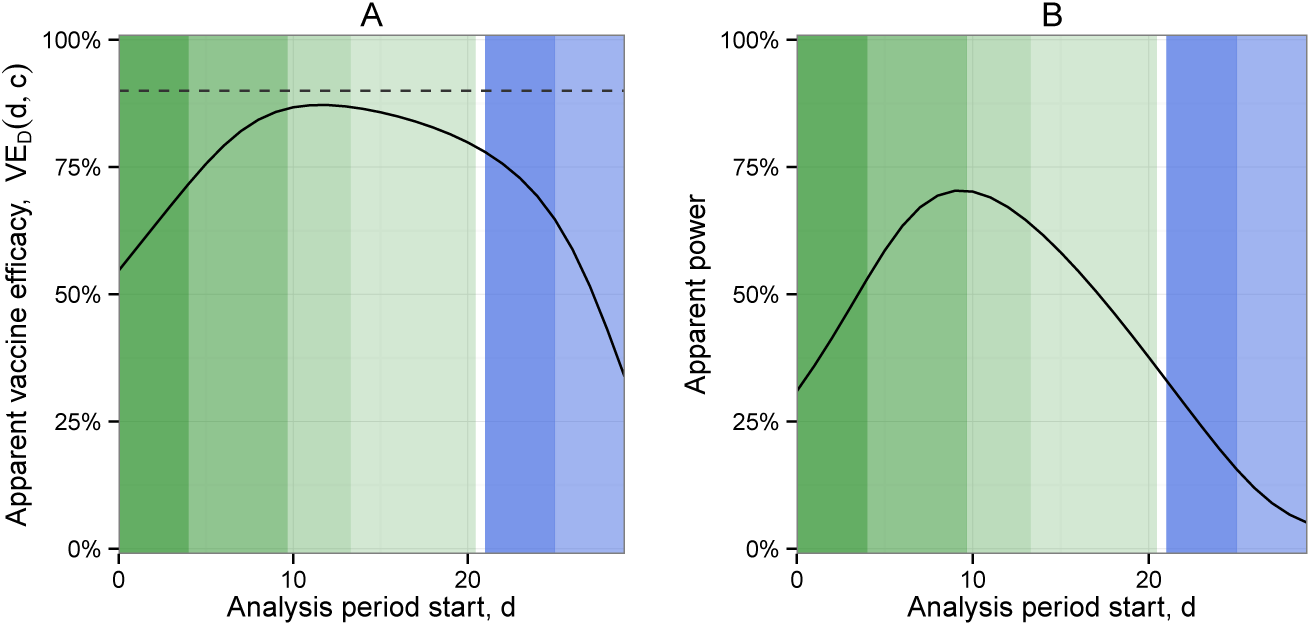
Arm 1 (immediate vaccination arm) vaccinated on day *s*_1_ = 0; Arm 0 (delayed vaccination) vaccinated on day *s*_1_ = 21 (*b* = 21). Constant background infection hazard rate *λ*_*W*_ = 0.001. Standard setting. Sample size *n* = 500 per arm. (Setting as in Figure 3.) Analysis period [*d, d* + 21) with width *c* = 21; a range of *d* values are considered. (A) Apparent VE to assess bias, where the dotted line indicates *VE*_0_. (B) Apparent power.

A natural way to decrease bias without negatively impacting power is to increase the length of the vaccination delay *b*. In Figure 7, we increase the vaccination delay to *b* = 35 days while maintaining an analysis period of width *c* = 21. By increasing the vaccination delay, there is a longer period of time in which Arm 1 (immediate arm) is protected and Arm 0 (delayed arm) is not protected. Thus, there is more observation time in which the apparent *VE*_*T*_ (*t*) is equal to the true *VE*_*S*_. As a result, we note that it is possible to identify a 21 day analysis period in which there is little to no bias (panel D), and bias and power are less sensitive to the choice of *d* (panels D and E). This stable region occurs roughly for *d* values between *R* + *u*_0.90_ to *R* + *u*_0.999_. This has the advantage that you have more tolerance for error when pre-specifying the delay. Note although in panel C the stable region extends slightly beyond 35 days, a 21 day analysis period would need to start well before then to limit bias in apparent VE, as shown in panel D. As described above, a more powerful approach would be to similarly increase the width of the analysis period *c*; this induces bias, though less bias than when *b* = 21 because *VE*_*T*_ (*t*) = *VE*_0_ for a larger proportion of the analysis period.

**Figure 7:**
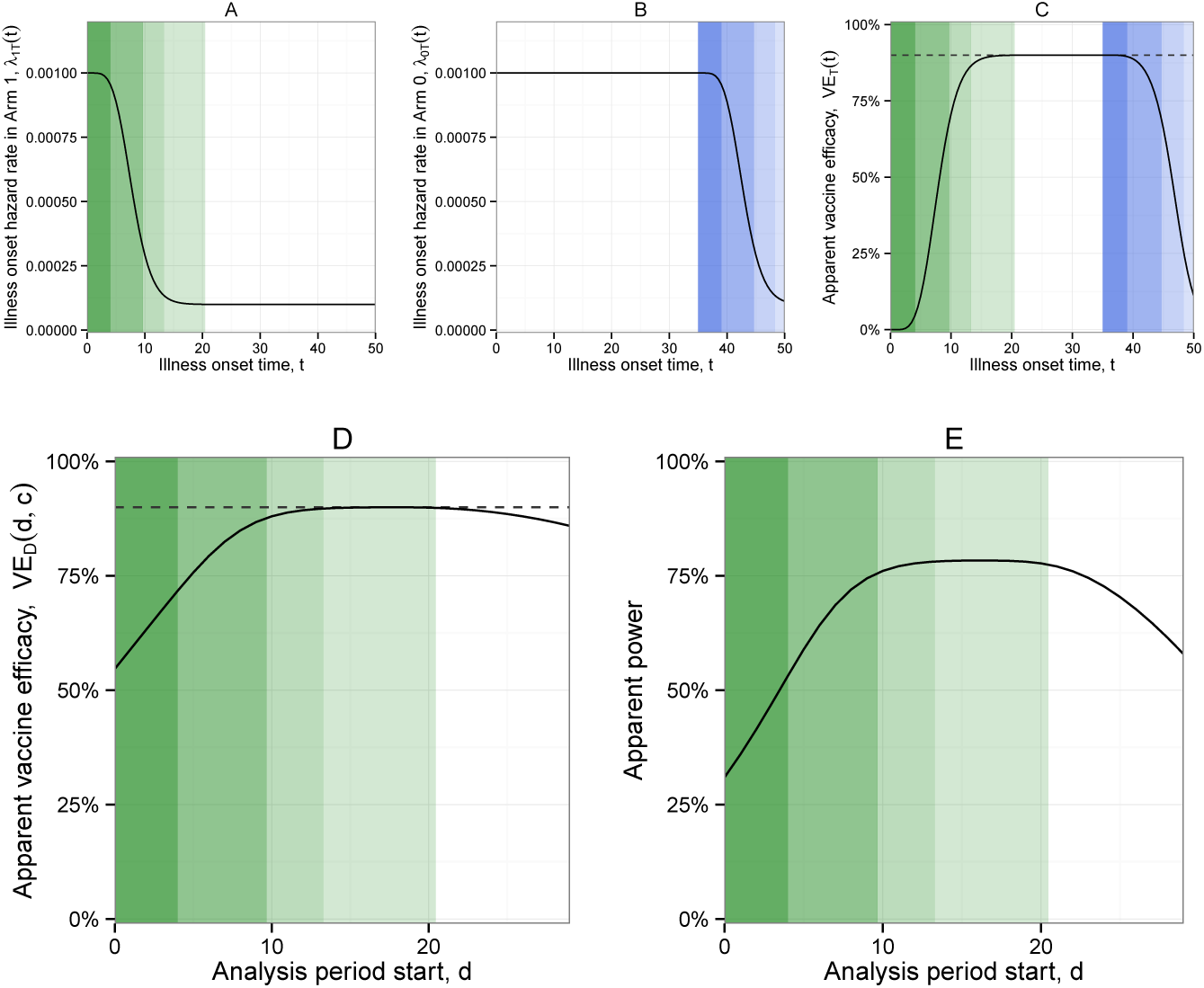
Arm 1 (immediate vaccination arm) vaccinated on day *s*_1_ = 0; Arm 0 (delayed vaccination) vaccinated on day *s*_1_ = 35 (*b* = 35). Constant background infection hazard rate *λ*_*W*_ = 0.001. Standard setting. Sample size *n* = 500 per arm. (Setting as in Figure 3.) Analysis period [*d, d* + 21) with width *c* = 21; a range of *d* values are considered. (A) Illness onset hazard rate in Arm 1 as a function of time. (B) Illness onset hazard rate in Arm 0 as a function of time. (C) Apparent VE (1 - illness onset hazard ratio) as a function of time, where the dotted line indicates *VE*_0_. (D) Apparent VE to assess bias, where the dotted line indicates *VE*_0_. (E) Apparent power.

In general, if bias is the primary concern, we suggest selecting *c < b*. Furthermore, we do not recommend counting events past *b* + *R* + *u*_0.50_, or, more stringently, *b* + *u*_0.50_. In some settings there may be pushback against counting any events in Arm 0 following delayed vaccination; thus the analysis period end can be set at *b*. Selecting *d* = *R* + *u*_0.90_ should reasonably minimize bias, though *d* = *R* + *u*_0.999_ could be selected as a more stringent option. Note that power may be severely impacted if the most stringent options are applied (see Figures S23 and S24 in Supplementary Materials). These are recommended as starting values, but the approximations and/or simulations should be used to incorporate context-specific factors. If maximizing power is the primary concern, selecting *c* = *b* is recommended to capture the greatest number of events, and *d* = *R* + *u*_0.50_ is proposed as a starting value.

Particularly in the context of emerging infectious diseases, the disease and/or vaccine may not be well characterized when the trial protocol is developed. We want the success of the trial to be reasonably robust to the selected analysis period. One strategy is to increase *b* to the maximum value that is ethically acceptable; as is evident above, this has a number of statistical advantages. Another strategy is to avoid trial designs for which the bias and/or power have narrow peaks at certain values of *d*, and recall that sometimes these are not the same values for bias and for power. A specific approach we recommend is to identify “plausibility ranges” for the unknown parameters and identify a trial design (combination of *d* and *c*) that has minimal bias for all combinations within these ranges; then, we suggest increasing the sample size until the desired power is achieved for that design.

## 7 Ebola vaccine trial example

The interim analysis of the ‘Ebola ¸ca suffit’ ring vaccination trial in Guinea provides a real application of the principles described above [Henao-Restrepo et al., 2015]. In this trial, a delayed vaccination arm was used with vaccination occurring 21 days after randomization as compared to an arm vaccinated immediately after randomization. The analysis period start was set at *d* = 10 days, excluding cases with illness onset between 0 and 9 days after randomization from the per protocol analysis. As no cases were observed in vaccinees in either arm more than 6 days after vaccination, the estimated VE was 100% (95% CI: 75.1, 100%). In fact, the estimated VE is largely insensitive to the choice of *d*, remaining 100% for any *d* after the last case in an immediately vaccinated individual. Similarly, the estimated VE is insensitive to the analysis period end, which is unique to the setting of *VE*_0_ = 100% (see Figure S25 in Supplementary Materials); because no further cases accumulate in either arm after a certain point, the point estimate does not change. As this paper demonstrates, a risk of delayed vaccination designs is bias towards the null; with VE of 100%, this type of bias was not observed in the trial. The power, on the other hand, is sensitive to the choice of *d*, with earlier values of *d* yielding greater power because more events in the delayed arm are retained.

## 8 Discussion

In this paper we present a framework for selecting the per protocol analysis period for vaccine efficacy trials. This framework accounts for the facts that only illness (symptom) onset times are observed, and illness onsets shortly after vaccination may reflect infections before vaccination or before the immune system developed protection. A sensitive baseline test, if available, can help identify infections occurring prior to vaccination, but it cannot identify infections occurring before the immune response has fully developed. The per protocol analysis seeks to exclude these early cases, along with participants who do not follow trial protocol, to achieve an unbiased estimate of vaccine efficacy. An unbiased estimate of vaccine efficacy is critical for deciding whether the vaccine product should be licensed, and delayed vaccination trials typically return an attenuated vaccine efficacy estimate. We provide closed-form approximations for predicting the bias and power for a given analysis period, and we describe the bias/variance tradeoff associated with counting these early cases.

Delayed vaccination can be used as a comparator arm in settings where placebo or vaccination for an unrelated disease is not considered ethically acceptable. This approach may be preferred for diseases with high case fatality rates, such as Ebola virus disease. The design also places the vaccine in areas of greatest need, potentially averting additional cases and enhancing disease control efforts if the vaccine is efficacious. Using a delayed vaccination comparator necessarily decreases study power as it limits the time when the two arms can be meaningfully contrasted for estimation of vaccine efficacy. Once delayed vaccination participants are protected by vaccine, the analysis must stop because no appropriate control remains. Selecting the appropriate analysis period is also crucial for limiting bias. Other challenges of the approach include that blinding may be difficult to achieve. If availability of vaccine is a limiting factor, it is noteworthy that delayed vaccination trials require more doses as participants in both arms are vaccinated. Finally, if the vaccine candidate requires multiple doses or takes a long time to develop efficacy, the delay would need to be very long to achieve the desired power, thereby reducing the ethical advantage of the approach.

The framework we provide is intended to support those designing vaccine efficacy trials in calculating sample size, power, and pre-specifying the appropriate analysis period. As for any sample size/power calculation, this approach requires a number of assumptions for which limited information may be available. The closed-form approximations had small discrepancies from simulation results, especially when the sample size was small (*<*500 per arm). The underlying model assumes independence and does not reflect the complicated nature of infectious disease dynamics, including indirect vaccine effectiveness. We suggest using the closed-form approximations to narrow down the space of trial designs while planning and using more realistic simulations as a confirmatory step. Other limitations include that the ramp-up period is assumed fixed and everyone is assumed to be vaccinated on the same day; these could be readily modified by converting these constants to random variables.

Immunological data from early phase trials can be used to support the design of a vaccine efficacy trial. If these data are rapidly accumulating, as in an emergency setting, and is not available at the time the protocol is written, the statistical analysis plan could include a clause allowing flexibility in the specification of *d* pending external data. Alternatively, a different value of *d* could be pre-specified as a secondary analysis. More sophisticated approaches could be used than applying hard cutpoints for the analysis period, though typically simpler approaches are preferred for clinical trial primary endpoints. The incubation and immune ramp-up periods could be explicitly included in the likelihood; if this lag is known, it can be used to construct optimal weights [Zucker and Lakatos, 1990]. Furthermore, we always recommend reporting survival (or cumulative incidence) curves so that any evidence of an early harmful effect is not missed [Hern´an, 2013].

Moving forward, the international community has recognized the need for specific guidance on how to design vaccine trials in emergency settings [World Health Organization]. For emerging pathogens with high case fatality rates, delayed vaccination may be a desirable strategy when evaluating vaccine candidates. This guidance is intended to support those implementing this approach.

## 9 Acknowledgments

National Institutes of Health grants U54-GM111274 and R37-AI032042, World Health Organization, Ana Maria Henao-Restrepo and the ‘Ebola ça Suffit’ trial consortium.

## References

S. E. Bellan, J. R. C. Pulliam, C. A. B. Pearson, D. Champredon, S. J. Fox, L. Skrip, A. P. Galvani, M. Gambhir, B. A. Lopman, T. C. Porco, L. A. Meyers, and J. Dushoff. Statistical power and validity of Ebola vaccine trials in Sierra Leone: A simulation study of trial design and analysis. The Lancet Infectious Diseases, 15(6):703–710, 2015.

A. Camacho, R. M. Eggo, S. Funk, C. H. Watson, A. J. Kucharski, and W. J. Edmunds. Estimating the probability of demonstrating vaccine efficacy in the declining Ebola epidemic: a Bayesian modelling approach. BMJ Open, 5(12):e009346, 2015.

Ebola ça Suffit Ring Vaccination Trial Consortium. The ring vaccination trial: a novel cluster randomised controlled trial design to evaluate vaccine efficacy and effectiveness during outbreaks, with special reference to Ebola. British Medical Journal, 351:h3740, 2015.

P. B. Gilbert, J. O. Berger, D. Stablein, S. Becker, M. Essex, S. M. Hammer, J. H. Kim, and V. G. DeGruttola. Statistical Interpretation of the RV144 HIV Vaccine Efficacy Trial in Thailand: A Case Study for Statistical Issues in Efficacy Trials. Journal of Infectious Diseases, 203(7):969–975, 2011.

M. E. Halloran, M. Haber, I. M. J. Longini, and C. J. Struchiner. Direct and Indirect Effects in Vaccine Efficacy and Effectiveness. American Journal of Epidemiology, 133(4):323–331, 1991.

A. M. Henao-Restrepo, I. M. Longini, M. Egger, N. E. Dean, W. J. Edmunds, A. Camacho, M. W. Carroll, M. Doumbia, B. Draguez, S. Duraff, G. Enwere, R. Grais, S. Gunther, S. Hossmann, M. K. Konde, S. Kone, E. Kuisma, M. M. Levine, S. Mandal, G. Norheim, X. Riveros, A. Soumah, S. Trelle, A. S. Vicari, C. H. Watson, S. Keita, M.-P. Kieny, and J.-A. Rottingen. Efficacy and effectiveness of an rVSV-vectored vaccine expressing Ebola surface glycoprotein: interim results from the Guinea ring vaccination cluster-randomised trial. The Lancet, 386(9996):857–866, 2015.

M. A. Hernán. The Hazards of Hazard Ratios. Epidemiology, 21(1):13–15, 2013.

A. D. Horne, P. A. Lachenbruch, and K. L. Goldenthal. Intent-to-treat analysis and preventive vaccine efficacy. Vaccine, 19(2-3):319–326, 2000.

M. A. Hussey and J. P. Hughes. Design and analysis of stepped wedge cluster randomized trials. Contemporary Clinical Trials, 28:182–191, 2007.

R Core Team. R: A Language and Environment for Statistical Computing, 2015. URL http://www.r-project.org/.

M. S. Ridout, C. G. Demétrio, and D. Firth. Estimating intraclass correlation for binary data. Biometrics, 55(1):137–148, 1999. ISSN 0006-341X.

B. Rosner. Fundamentals of Biostatistics. Cengage Learning, 7th edition, 2010. ISBN 978-0538733496.

RTSS Clinical Trials Partnership. Efficacy and safety of RTS,S/AS01 malaria vaccine with or without a booster dose in infants and children in Africa: final results of a phase 3, individually randomised, controlled trial. The Lancet, 386(9988):31–45, 2015.

T. M. Therneau. Package ’survival’: Survival Analysis. R package, 2015. URL http://cran.r-project.org/web/packages/survival/index.html.

World Health Organization. A research and development blueprint for action to prevent epidemics. URL http://www.who.int/csr/research-and-development/en/.

R. Xu and J. O’Quigley. Estimating average regression effect under non-proportional hazards. Biostatistics, 1(4):423–439, 2000.

D. M. Zucker and E. Lakatos. Weighted log rank type statistics for comparing survival curves when there is a time lag in the effectiveness of treatment. Biometrika, 77(4):853–864, 1990.

